# A preliminary description of the ecological characteristics of wild waterbird Japanese encephalitis virus hosts in high risk landscapes in India

**DOI:** 10.1101/2022.01.13.476136

**Authors:** Michael G. Walsh, Amrita Pattanaik, Navya Vyas, Deepak Saxena, Cameron Webb, Shailendra Sawleshwarkar, Chiranjay Mukhopadhyay

## Abstract

Wild reservoirs of Japanese encephalitis virus are under-studied globally, which presents critical knowledge gaps for JEV infection ecology despite decades of received wisdom regarding this high-impact mosquito-borne virus. As a result, ardeid birds, generally understood to be the primary reservoirs for JEV, as well as other waterbirds occupying landscapes at high risk for spillover to humans, are frequently ignored by current surveillance mechanisms and infrastructure. This is particularly true in India, which experiences a high annual burden of human outbreaks. Incorporating wild reservoirs into surveillance of human and livestock populations is therefore essential but will first require a data-driven approach to target individual host species. The current study sought to define a preliminary ecological profile of JEV hosts based on 1) species ecological traits, and 2) species presence and abundance adjusted for the biotic constraints of sympatry. Optimal host species tended to be generalists and demonstrate regionally-increasing populations. While ardeid bird species richness, abundance, and relative abundance did demonstrate the strongest and most consistent associations with the distribution of human JEV outbreaks, this study also identified several individual species among two other bird families in these landscapes, the Anatidae and the Rallidae, which also exhibited an optimal host profile and were strongly associated with the distribution of outbreaks. The findings from this work provide the first data-driven evidence base to inform wildlife sampling for the monitoring of JEV circulation in outbreak hotspots in India and thus identify good preliminary targets for the development of One Health wildlife JEV surveillance.

## Introduction

Japanese encephalitis virus (JEV) is a mosquito-borne zoonotic virus circulating enzootically in wild waterbirds and domestic pigs, and seasonally spills over to humans causing disease (Japanese encephalitis (JE)) with extensive morbidity and mortality, particularly in children[1,2]. India experiences high JEV incidence, with large outbreaks clustering in the northeast, and to a lesser extent in the southwest, during the monsoon season[3]. *Culex tritaeniorhynchus* is the primary JEV vector across Asia, and also plays a substantial role in transmission in India[4–6]. In addition to *Cx. tritaeniorhynchus*, there are four more vector species (*Cx. vishnui, Cx. gelidus, Cx.fuscocephala*, and *Cx. pseudovishnui*) widely distributed across India[6,7]. The near-ubiquitous presence of vectors notwithstanding, JEV outbreaks in humans manifest within distinct landscape mosaics. Landscapes of fragmented wetland-rainfed agricultural mosaics, and exhibiting an extensive distribution of ardeid birds and domestic pigs and chickens, presented the greatest risk of JEV outbreaks across India in a recent report of national JEV surveillance between 2010 and 2020 [8].

Despite the substantive annual burden of disease associated with JEV, there is a surprising dearth of knowledge regarding the infection ecology of this arbovirus, particularly with respect to its circulation in wildlife populations. Bird species in the Ardeidae family have long been recognised as key maintenance hosts of JEV[9–13], while domestic pigs have been implicated as the primary amplification and bridging hosts for human spillover[14–20], although additional evidence has shown that some ardeid species can also amplify JEV circulation sufficiently to facilitate direct spillover to humans via the mosquito vectors[4]. There is also some evidence to suggest that chickens may also play a role as amplifying hosts, further expanding the potentially relevant interspecific interactions at the wildlife-livestock interface[12,21,22]. Despite the accepted state of the knowledge regarding JEV reservoir hosts, much of the wildlife survey data that this is based on is outdated, limited to only a small number of ardeid species, and was not collected in India. The most extensive investigation of human outbreaks in India to date provided strong support for the association between JEV outbreaks and both ardeid species distributions and pig density[8]. Importantly, this study also identified mosaics of riparian and freshwater marsh wetlands with fragmented rainfed agriculture as the key landscapes in which these hosts contribute to JEV circulation and the high risk of outbreaks. However, no individual ardeid species associated with these landscapes were explored in this study. While JEV isolation from ardeids has been reported outside of India, despite several serological surveys, only one study has isolated virus from ardeid birds in India[13]. In addition, there are other waterbird species associated with such landscapes, some of which have also been identified as JEV hosts, but again all but one of these were based on serological surveys. A more thorough evaluation of the presence and ecological characteristics of individual waterbird species that may function as JEV hosts in these landscapes is therefore warranted in order to identify preliminary target species for the development of much-needed JEV surveillance that incorporates wildlife monitoring.

The study of the ecology of enzootic hosts has increased in recent years as a means to help classify potential hosts based on specific biological and ecological traits, frequently focusing on life history due to the association between fast-living and host competence identified in many r-selected species[23–29]. These species, which are typically generalist and frequently more resilient to anthropogenic pressure[30–32], tend to be fast-living and invest more in earlier and more fecund reproductive effort and less in innate immune function and adult survival[33]. As such, many of these species also present as optimal hosts for pathogens[34]. Nevertheless, the association between fast-living species and host competence is by no means universal, with some systems demonstrating the opposite or ambiguous associations, such as ebolaviruses[35], Rift Valley fever virus[36], and Ross River virus[37]. In addition to life history, other host traits, including population density, diet breadth, foraging strategy, and habitat disturbance, can also be influential to the infection ecology of some pathogens[31,32]. Japanese encephalitis virus host ecology has hitherto gone unexplored among ardeid, or other waterbird, species within India and thus represents a critical knowledge gap for this system.

The objectives of the current study were three-fold. First, we sought to define an ecological trait-based JEV host profile for the wild waterbird species that occupy the landscapes of high risk for JEV spillover in India. Second, we estimated the species landscape suitability of each of these bird species, as well as species richness, to compare whether the distributions of hosts identified by the trait-based approach are associated with the distribution of JEV outbreaks. Third, we further adjusted the presence and abundance of each bird species, as well as overall and family-specific species richness, for the potential biotic constraints of sympatry at the community level and again compared these adjusted metrics to the distribution of JEV outbreaks to identify optimal target species for wildlife surveillance in high-risk landscapes at the community level.

## Methods

### Data sources

Seventy-six species in the families Ardeidae, the herons (including egrets) and bitterns (n = 19), Anatidae, the ducks, geese, and swans (n = 41), and Rallidae, the rails, coots, crakes, and gallinules (n = 16) are documented as extant in India[38]. These families comprise the waterbird species that occupy the wetland habitat previously identified as high risk for JEV outbreaks[8]. Each of these families has at least one species with documented susceptibility to JEV infection. For the purposes of qualifying JEV infection in waterbird hosts below, infection competence is defined here as a viral titre in a host species of sufficient magnitude to infect vector mosquitoes and thus pass on the infection to other hosts[26]. Infection classification by serology only can designate a host’s susceptibility to infection, but not competence. The phylogenetic tree of these species along with the distribution of each family across India is presented in Figure 1. The phylogenetic tree was obtained from the VertLife project[39] and constructed based on the methods described by Jetz et al.[40]. Any taxonomic binomial discrepancies between the VertLife tree and the named species obtained from the Global Biodiversity Information Facility used for the species distribution models described below were resolved, with alternate binomials appearing in parentheses below. Among the Ardeidae, *Egretta garzetta*[10,41,21], *Areola grayi*[9,10,13], *Nycticorax nycticorax*[11,41,21], *Mesophoyx intermedia* (*Egretta intermedia*)[11,41,21], *Casmerodius albus* (*Ardea alba*)[42], *Bubulcus ibis* (*Bubulcus coromandus*)[9,42], have documented JEV infection, all with demonstrated infection competence. Among the Anatidae, *Anas platyrynchos*[42,43], *A. poecilorhyncha*[43], *A. crecca*[43], *A. acuta*[43], *A. strepera*[44], *A. penelope*[43], *A. clypeata*[44], *A. formosa*[43], *Aythya fuligula*[44], and *Aix galericulata*[43], have documented JEV infection, but only one (*A. platyrynchos*) with demonstrated infection competence[42]. Infection susceptibility in the remaining anatid species was documented by serology only. Among the Rallidae, *Gallinula chloropus*[9] and *Fulica* atra[43] have documented JEV infection by serology, but neither have documented infection competence. These 18 species were classified as JEV susceptibility-positive, while the remaining species in these three families were classified as undetermined JEV status. Because multiple testing modalities (e.g. serology vs. virus isolation) were employed in the field surveys describing these species’ infections, it must be emphasised that none of these species can be designated as reservoir, maintenance, or amplification hosts in the comparison that follows below. Host species are designated only as susceptible to JEV infection and we concede the attendant limitations to inference based on this classification. Furthermore, potential bias may be introduced due to differences in reporting effort across different bird species. To address this potential bias, reporting effort was quantified using the total number and the number of India-specific published studies for each species in the Web of Science (WoS) database. This metric for reporting effort was then included as a covariate in subsequent models (see statistical analysis below), following the approach of previous work[45,46].

**Figure 1.**
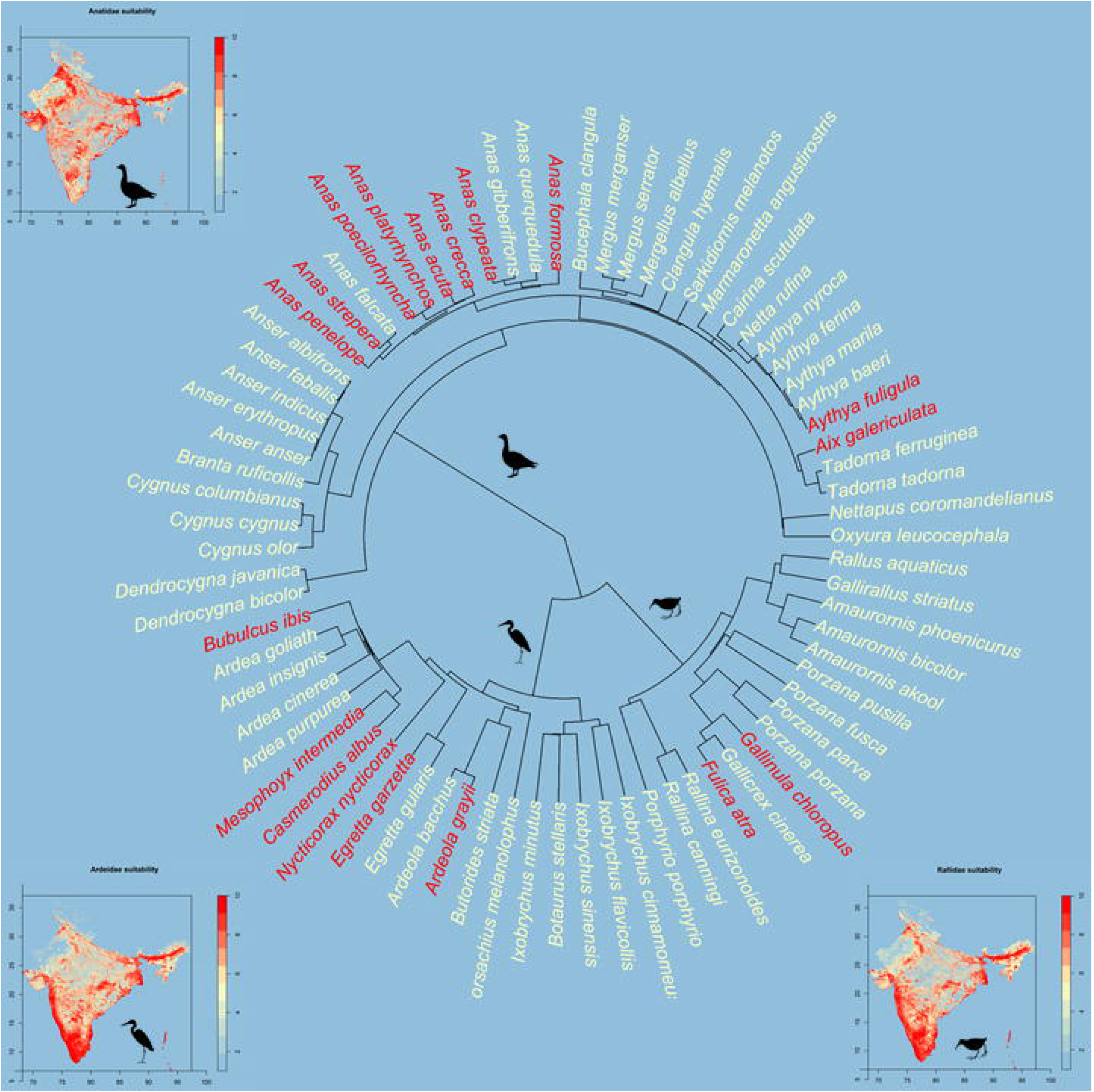
The phylogenetic tree of the waterbird species that occupy high risk Japanese encephalitis virus landscapes with the distribution of species richness within each family presented peripherally to the tree. Family taxa are presented in silhouette. Maps are used only for the purposes of species richness and do not reflect the authors’ assertion of territory or borders of any sovereign country including India.

Bird species’ diet composition, foraging strategy, and body mass were obtained from the Elton Traits database[47]. Diet composition and breadth may be important characteristics of host species insofar as these can delineate generalist versus specialist species and potential resilience to human pressure[48], whereas body mass is an important life history characteristic as well as being a potential modulator of immune function[26,49]. Three broad dietary classifications were compared. First, the proportion of the diet derived from plants was aggregated across all potential plant sources (seeds, fruit, nectar, grasses, leaves, and other plant material) to characterise the overall proportion of the diet derived from plants. Second, the proportion of the diet derived from vertebrates was aggregated across all potential vertebrate food sources (fish, mammals, birds, reptiles, and amphibians). Third, the proportion of the diet derived from invertebrates was similarly classified. In addition, binary classifications of diet strategy comprised 1) frugivorous/nectivorous, 2) granivorous/herbivorous, 3) invertivorous, 4) vertebrate-eating, and 5) omnivorous. The foraging strategies considered here comprised six categories: 1) foraging below the water surface at depth, 2) foraging at or around the water surface, or in shallows less than 12.7 cm, 3) foraging on dry ground, 4) foraging in understory growth less than 2 m in height, 5) foraging in middle-story growth greater than 2 m in height but below the tree canopy, and 6) foraging in the tree canopy[47].

The TetraDENSITY database was used to obtain species mean population density[50], while the reproductive characteristics, clutch size and egg mass, were acquired from the Lislevand bird traits database[51]. These species traits were included since population density may influence the dynamics of JEV circulation in populations and communities[52–55], and clutch size and egg mass are primary indicators of reproductive life history. The data on population density, clutch size, and egg mass were missing for 50%, 38%, and 25%, respectively, so a random forest machine learning algorithm, which was previously shown to be a robust approach to this application[25,56], was used to impute the missing data. The rflmpute function in the randomForest package was used to implement the algorithm[57].

The International Union for the Conservation of Nature (IUCN) Red List was used as a source of several landscape features associated with each bird species[58]. The IUCN variables were employed as a means of describing the relevance of anthropogenic pressure to species’ JEV infection susceptibility status. First, the IUCN threatened status was obtained for each species. Species designated status was based on the following categories from greatest to least threatened: “extinct in the wild”, “critically endangered”, “endangered”, “vulnerable”, “near threatened”, and “least concern”. Species with IUCN designation “vulnerable”, “endangered”, or “critically endangered” were re-classified as “threatened”, while species with designation “least concern” or “near threatened” status were re-classified as “non-threatened” in a newly created dichotomous IUCN threat classification variable. No species included in the study were classified as “extinct in the wild”. Second, well-delineated threats due to logging, agricultural development, urbanisation, extractive industries, road/rail corridor development, climate change, hunting or harvesting activities, or incursion by invasive species were recorded as present or absent for each species. The association between each threat and JEV status was assessed individually, as well as combined over all threats in aggregate. Third, species population growth was recorded as increasing, decreasing, stable, or unknown.

All observations of Ardeidae species (1,016,733 individual observations of 16 species[59]), Anatidae species (494,863 individual observations of 28 species[60]), and Rallidae species (377,672 individual observations of 15 species[61]) between 1 January 2010 and 31 December 2020 across India were acquired from the Global Biodiversity Information Facility (GBIF) to model each species’ distribution, as well as overall species richness and species richness within family. After removing duplicate observations at the same geographic location and those species with an insufficient number of observations available for modelling (n < 100), there remained 241,784 observations of 15 Ardeidae species, 113,427 observations of 21 Anatidae species, and 78,372 observations of 12 Rallidae species for the landscape suitability models described below. *Bubulcus coromandus* was until recently considered a subspecies of *B. ibis* (*B. ibis coromandus*), and is represented in the GBIF database as both *B. ibis* and *B. coromandus*.

Because of the potential for differential accessibility, and thus differential reporting of bird occurrence, the background points used to model all species distributions were selected proportional to the human footprint (HFP) (see modelling description below) as a proxy for accessibility, thus correcting for potential spatial reporting bias. The HFP raster was obtained from the Socioeconomic Data and Applications Center (SEDAC) registry[62] and quantified according to a 2-stage classification system that has been described in detail[63]. Briefly, a metric for human influence was first quantified based on human population density, rural versus urban location, land cover, degree of nigh time light pollution, and proximity to roads, rail lines, navigable rivers, and coastline. The domains were scored and summed to generate the human influence index (HII), and then HFP was calculated as the ratio of the range of minimum and maximum HII in the local terrestrial biome to the range of minimum and maximum HII across all biomes, expressed as a percentage[63].

Two-hundred and ninety-four laboratory-confirmed outbreaks of JEV were reported to the National Centre for Disease Control’s Integrated Disease Surveillance Programme (IDSP) at a spatial resolution of 1 arc minute between 1 January, 2010 and 31 December, 2020 and have been described previously[8]. The IDSP maintains a national JEV surveillance system under the administration of India’s Ministry of Health and Family Welfare[64], These surveillance data were also previously externally-validated against an independent, laboratory-confirmed dataset of community surveys of human and mosquito infection[8].

Communities of low socioeconomic status are disproportionately affected by JEV outbreaks. Since these communities frequently experience limited access to health care, the current study used the distribution of health system performance as a proxy for the local capacity to detect cases and thereby adjust for potential reporting bias of JEV infections (see modelling description below). The infant mortality ratio (IMR) was chosen as the best metric for health system performance because it has been validated as a robust measure of health infrastructure and health system performance and is frequently used to evaluate health service delivery and performance across heterogeneous socio-geography [65,66]. The IMR is strongly correlated with the Human Development Index (HDI) and the Inequality-Adjusted Human Development Index (IHDI) and is considered a reasonable representation of the environmental, economic, and social structural determinants of population health[65,67]. The IMR raster was obtained from the SEDAC repository[68].

The WorldClim Global Climate database was the source of the climate data used in this study[69]. These comprised mean annual temperature, mean annual precipitation, and isothermality. Proximity to surface water was calculated using the proximity function in the QGIS geographic information system[70] and subsequently used to generate a distance raster for the hydrogeography data obtained in the Global Lakes and Wetlands Database[71,72], The pixel values of this raster represent the distance in kilometres between surface water and all other pixels within the geographic extent of India.

### Statistical Analysis

#### Species trait analysis

Phylogenetic generalized linear models (PGLMs) fit with the binomial family were used to evaluate the associations between species’ ecological traits and their JEV status while accounting for the phylogenetic correlation across species[73]. As described above, the degree of “studiedness” of each species, globally and within India, was determined using WOS citations and included as a covariate to correct for potential bias introduced by differences in reporting effort across species. Twenty-six bivariate models were fitted wherein each model comprised a simple PGLM assessing the crude association between JEV susceptibility-status and each species trait: reporting effort (global), reporting effort (India-specific), body mass, clutch size, egg mass, population dynamics, threatened due to agriculture, urbanisation, logging, extractive industries, road/railway development, hunting, invasive species, or climate change, total number of anthropogenic threats, dietary categories, IUCN threat status, forage strategies, and population density. Those trait features associated with JEV status were then selected and fitted together to assess a multivariable PGLM. The ape package[74,75] in the R statistical environment[76] was used to quantify the phylogenetic correlation, and the phylolm package was used to fit the PGLMs[77].

#### Species distribution modelling

There were 48 of the 76 described extant species in India with a minimum of 100 unique observations available in the GBIF database (S1 Table 1). This was the minimum number of observations we considered appropriate to estimate the landscape suitability of each species. Landscape suitability was estimated using an ensemble approach comprising boosted regression trees (BRT), random forests (RF), and generalised additive models (GAM). Boosted regression trees and RF both partition data space by optimising homogeneity among predictors and a response (i.e. species presence). The algorithms generate and combine many decision trees, resulting in optimised decision trees that reduce overfitting and can capture complex interactions between the predictors[78–81]. There are two important differences between BRT and RF, however. With RF, only a random subset of predictors is selected from the set of all predictors for the generation of each decision tree. This reduces overfitting by decorrelating the data through the random selection of predictors for each tree. With BRT, overfitting is reduced by growing trees sequentially and learning from the previous iteration rather than decorrelating trees based on the sampling of subsets as with RF. In contrast, the GAM framework fits multiple basis functions for smoothed covariates thus allowing for nonlinear relationships between outcomes and covariates[82,83]. Each model under BRT, RF and GAM was fit with five-fold crossvalidation. Observation data were thinned so that only one observation per pixel was included in the analysis to avoid artificial spatial clustering (S1 Table 1). Mean annual temperature, mean annual precipitation, isothermality, and proximity to surface water were included as landscape features at 30 arc sec resolution in all models. For each species, each of the three models (BRT, RF, and GAM) fitted to their occurrence and background points was evaluated by model performance, based on the area under the receiver operating characteristic curve (AUC), and model fit, based on the deviance. Landscape suitability for each species was then derived from an ensemble of the three models (BRT, RF, and GAM) using their weighted mean with weights based on AUC[84]. To correct for potential spatial sampling bias among the GBIF observations, background points were sampled proportional to the human footprint to serve as a proxy for landscape, and thus bird, accessibility. Each species’ landscape suitability as derived from the ensembles was subsequently summed across all species as an estimate of local species richness across the geographic extent of India. This estimate was then adjusted further for the biotic constraints of sympatric species at the community level (see *Community-level modelling* description below).The sdm package[84] was used for fitting all models and deriving the landscape suitability ensembles.

#### Point process modelling

The individual species distributions of those species predicted to be optimal hosts by the PGLM were tested to determine if they were associated with JEV outbreaks by fitting inhomogeneous Poisson point process models (PPMs) to those outbreaks[85,86]. This allowed the spatial dependence of the outbreaks’ distribution to be evaluated specifically with respect to each species’ landscape suitability. To control for potential reporting bias in outbreak surveillance, the background points used in these models were sampled proportional to IMR, as described above. A simple bivariate PPM of JEV outbreaks was thus created for each species identified as optimal by the PGLM, as well as any additional species previously documented as susceptible to infection but not otherwise flagged as an optimal potential host by the PGLM model. All PPMs were aggregated up to 1 arc minute, which was the scale of outbreak reporting[8]. The Akaike information criterion (AIC) was used to evaluate model fit and the AUC was used to assess model performance using an independent, laboratory-confirmed set of infection data, as described above and previously[8]. The spatstat R package was used to fit the PPMs[87].

#### Community-level modelling

To further adjust species richness for the potential interaction between sympatric species, and to compute individual species presence and abundance, interspecific biotic constraints were applied at the scale of the taluk. Taluks are 3rd-level, subdistrict municipalities that are sufficiently small to reasonably approximate shared space among sympatric species, but which are also sufficiently large enough to demarcate the minimal municipal infrastructure required across most of India for organising and executing animal and human disease surveillance. The biotic constraints took the form of sympatric species adjustments to the estimates of each species landscape suitability using the spatially-explicit species assemblage modelling (SESAM) framework[88–90]. First, the landscape suitability for each species was estimated using the ensemble method described above. Second, the individual species distributions were summed to calculate an unadjusted species richness estimate. Third, each species distribution is evaluated with respect to all other species present within each taluk via the probability ranking rule[89,91] to determine whether a given species should be retained within, or excluded from, each taluk “community”. Under this final step, each species is ranked from highest to lowest based on their suitability estimate obtained in the first step. Those with high suitability are ranked high, while those with low suitability are ranked low. Species are then selected for inclusion in the community starting with the species with the highest suitability estimate and continuing down through the list of ranked species until the sum of selected species is equal to the expected species richness value for each location as represented by the calculation in the second step. Once this threshold sum is reached, the adjusted-species richness is achieved and no further species are included in that particular taluk community. This process thus yields an estimate of individual species presence within taluks given the potential biotic constraints of the other species within those same taluks. Under this SESAM framework, for species retained as present following adjustment for taluk-level sympatric species, each 1 km^2^ pixel of their ensemble landscape suitability estimate raster that was greater than or equal to the true skill statistic (TSS)[92] was classified as present (1 = present, 0 = otherwise) and summed across all pixels within the taluk. In this way taluk-level species abundance was estimated for each species retained in a given taluk. The relative abundance of each species was then calculated by dividing each individual species abundance by the total abundance of all species retained as present for each taluk. New estimates of species richness for the Ardeidae, Anatidae, and Rallidae families were also generated by summing the number of species retained within each taluk, for each family, under the SESAM framework. As such this approach provided a framework for taluk-level community estimation of individual species’ abundance, as well as species richness, all of which were adjusted for sympatric species. The SESAM analysis was conducted in R using the ecospat package[90].

Similar to the local point process estimation described above, the taluk-level abundance of those species predicted to be optimal hosts by the PGLM, adjusted for the biotic constraints of sympatric species, were interrogated to determine if associations with JEV outbreaks were maintained having accounted for community sympatry. Integrated nested Laplace approximation (INLA) models[93] were used to estimate these biotically constrained associations at the community level of the taluk. It is important to note that any such associations identified do not provide any specific insight into species’ roles as hosts since species competence for JEV was not measured or evaluated in the current study. Individual species associations were instead explored simply to identify which species may be optimal as sampling targets for implementing new wildlife surveillance in outbreak hotspots across India, or as sentinels of JEV outbreaks. The INLA models were fit using the binomial likelihood family and with Besag–York–Mollie priors for the random effects. Taluks were modelled as either outbreak-positive or outbreak-negative over the duration of the study, and were thus fit with the binomial family, since most of the taluks across India did not experience outbreaks, while those that did only experienced one or a very small number of outbreaks. The Watanabe-Akaike information criterion (WAIC) was used to assess the fit of all INLA models. The INLA package in R[93] was used to fit the INLA models.

## Results

The phylogenetic tree of extant Ardeidae, Anatidae, and Rallidae species in India is presented in Figure 1 along with the distribution of estimates of species richness within these families across the country. Boxplots of ecological traits by JEV status are presented in S2 Figure 1, and individual simple PGLMs for bivariate associations between each trait and JEV status are presented in S3 Table 2. The final PGLM is presented in Table 1. Species spending a greater proportion of their time foraging at the water’s surface or in the shallows, species with increasing population growth, and omnivorous species were all more likely to be classified as optimal JEV host targets. The model performed well, with the AUC equal to 81%. Thirteen of the 23 interrogated species were positively associated with the distribution of JEV outbreaks at fine scale in the PPMs unadjusted for sympatry (S4 Table 3).

**Table 1.** Multiple phylogenetic generalised linear model presenting the associations between Japanese encephalitis hosts status and species ecological traits. Model fit was assessed using the Akaike information criterion (AIC) and model performance was assessed using the area under the receiver operating characteristic curve (AUC).

| Species trait | Regression coefficient | 95% confidence interval | p-value |
| --- | --- | --- | --- |
| Forage strategy – around water surface | 0.03 | 0.01 – 0.05 | 0.002 |
| Diet – omnivorous | 2.12 | 0.53 – 3.84 | 0.009 |
| Increasing population | 1.60 | 0.15 – 3.05 | 0.03 |

AIC = 89.62; AUC = 0.81

When sympatric species at the taluk level were accounted for in the estimation of each species’ presence and abundance a narrower range of optimal JEV host targets was identified (S5 Table 4). The species abundance of 3 ardeids (*B. coromandus, Butorides striata*, and *E. intermedia*), 5 anatids (*A. penelope, A. platyrhynchos, A. strepera, A.fuligula, Tadorna tadorna*), and 1 rallid (*Amaurornis akool*) were positively associated with the distribution of JEV outbreaks (Figure 2). Although optimal JEV hosts were identified in each of the three families, sympatry-adjusted within-family species richness, abundance, and relative abundance, were most strongly associated with the Ardeidae family, further supporting this family’s dominance in high-risk landscapes (Table 2 and S6 Figure 2).

**Figure 2.**
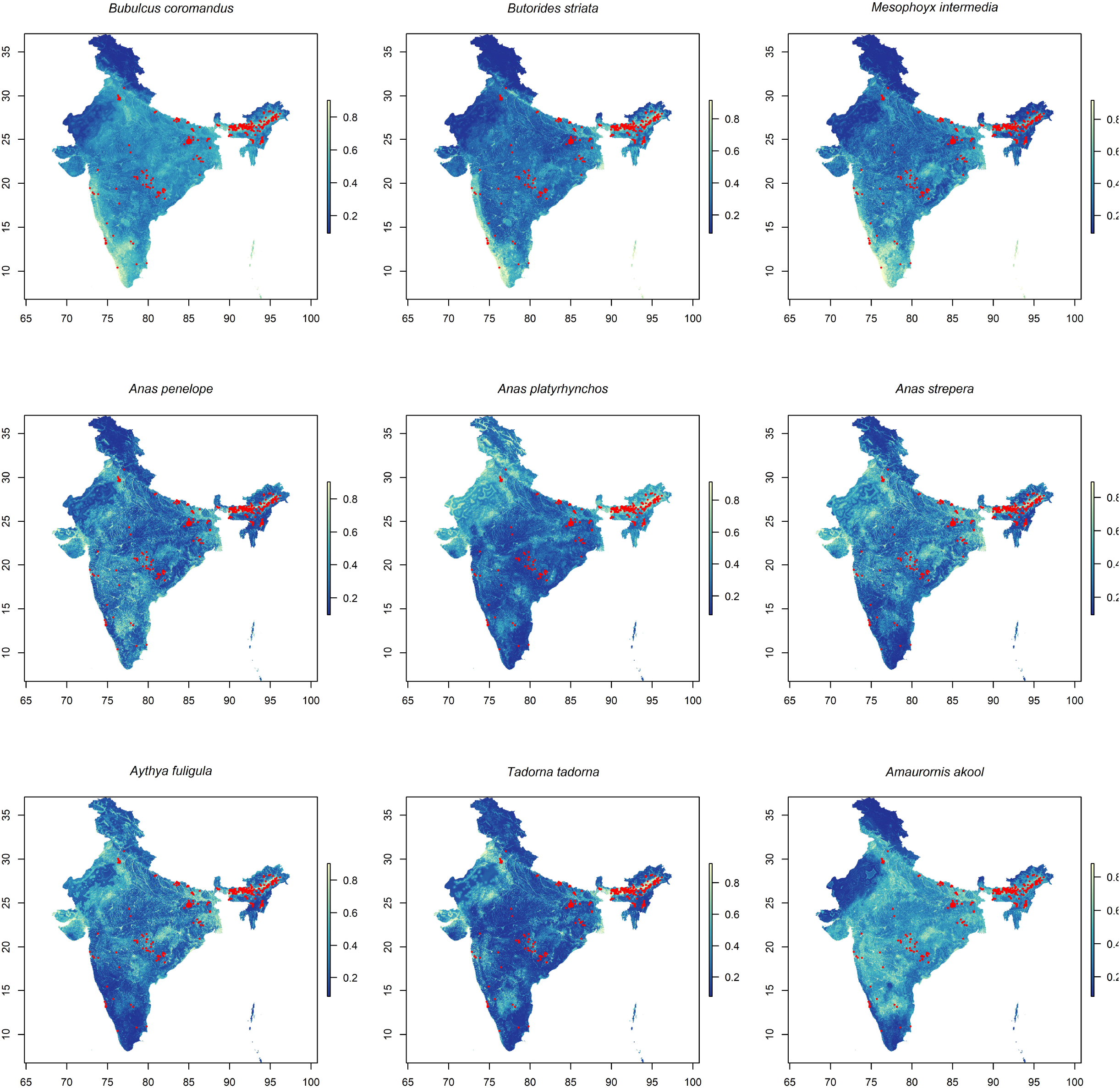
The distribution of the landscape suitability of those waterbird species identified as optimal potential targets for wildlife surveillance based on corroboration between trait-based phylogenetic generalised linear models and integrated nested Laplace approximation models after adjusting species presence for the biotic constraints of sympatric species. The overlaid red points represent the location of human JEV outbreaks. Maps are used only for the purposes of species richness and do not reflect the authors’ assertion of territory or borders of any sovereign country including India.

**Table 2.** Taluk-level integrated nested Laplace approximation models of Japanese virus outbreaks. Within-family species richness, family total abundance, and family relative abundance were all adjusted for the biotic constraints of sympatric species using the SESAM framework. Model fit was assessed using the Watanabe-Akaike information criterion (WAIC).

| **Taluk-level bird community features** | **Coefficient** | **95% credible interval** | **WAIC** |
| --- | --- | --- | --- |
| **Ardeidae** |  |  |  |
| Within-family species richness | 0.161 | 0.051 – 0.272 | 567.16 |
| Family total abundance | 0.0001 | 0.00004 – 0.0002 | 564.58 |
| Family relative abundance | 3.003 | 0.913 – 5.191 | 562.36 |
| **Anatidae** |  |  |  |
| Within-family species richness | 0.039 | -0.046 – 0.123 | 571.92 |
| Family total abundance | 0.000057 | 0.000026 – 0.00009 | 564.73 |
| Family relative abundance | 1.556 | -0.055 – 3.224 | 570.83 |
| **Rallidae** |  |  |  |
| Within-family species richness | 0.111 | -0.081 – 0.306 | 568.65 |
| Family total abundance | 0.00016 | 0.000065 – 0.00026 | 563.94 |
| Family relative abundance | -2.895 | -4.640 – -1.240 | 568.21 |
| **All birds** |  |  |  |
| Total species richness | 0.070 | 0.012 – 0.127 | 569.70 |
| Total species abundance | 0.00004 | 0.00002 – 0.000066 | 561.43 |

## Discussion

This is the first evaluation of population and community ecology characteristics of potential wild waterbird JEV hosts across India, the results of which can be used to identify initial targets for wildlife surveillance of this important zoonotic arbovirus. Several important features were in evidence. First, optimal hosts tended to be generalist species with regional population dynamics suggestive of growth. Second, this work provides the first country-wide, community-level estimates of water bird species richness and abundance in freshwater wetlands across India and adjusted for sympatry. These estimates were used to corroborate the predictions of optimal potential host species based on the phylogenetic trait-based modelling described above. While reaffirming the overall dominance of Ardeidae species relative to Anatidae and Rallidae species, this synthesis nevertheless identified several species across all three families that should provide good preliminary targets for the development of improved One Health wildlife JEV surveillance.

Animal surveillance for JEV in India is minimal in general, and virtually non-existent in wild birds in particular. Because the circulation of JEV in these hosts is fundamental to the virus’ infection ecology, there is a critical need to develop and implement surveillance infrastructure that incorporates the monitoring of reservoir birds in landscapes at high risk for JEV outbreaks, which are delineated by mosaics of wetlands and rainfed agriculture throughout India[8]. Unfortunately, given the limited field investigations to inform the selection of optimal targets for waterbird monitoring, a recognised and reliable source of known hosts from which to sample does not currently exist and so any development of wildlife monitoring for JEV must proceed with a minimal evidence base. In order to provide a more evidence-based list of potential JEV hosts for preliminary surveys, the current study interrogated the waterbird families of high-risk landscapes in India using trait-based population and community ecology methods, which allowed for an accounting of phylogenetic correlation and sympatry among species. This should provide a more sound approach than simply sampling birds previously identified as infected, most of which reflect surveys conducted outside of India, many years ago, or relied upon serology alone. Critical to determining the optimal species for initial sampling was the corroboration of species’ distributions with JEV outbreaks across India. Several of the species identified as optimal for sampling, including *B. coromandus, E. intermedia*, and *A. platyrhynchos*, have been previously demonstrated as competent hosts[41,21,42], and therefore reinforce these species as good targets for initial sampling and wildlife monitoring. The value of these results for the development of new wildlife surveillance notwithstanding, it is important to emphasise that these findings should not be interpreted as identifying any species as definitive maintenance, amplification, or bridging hosts. Host competence was not assessed in the current study and therefore the findings should not be used to delineate the infection ecology of JEV. Rather, this work is intended to inform sampling strategies for the implementation of field investigation and broader wildlife surveillance infrastructure, which is largely absent across India and which will be critical to understanding the circulation of JEV in wild waterbirds and thereby inform the landscape epidemiology of JEV outbreaks in humans. Interestingly, this study supported the dominance of ardeid bird presence in landscapes at high risk of JEV outbreaks relative to anatid and rallid bird presence, but also indicated that some species in these other waterbird families should also be considered for targeted surveillance sampling in these landscapes.

In addition to the important caution against over-interpretation of the results due to the inherent limitations described above, some additional limitations warrant further discussion here. First, there are a limited number of species included in this study with documented infection with JEV, largely due to the limited number of field surveys investigating JEV hosts in species extant in India. As such, potential bias due to differential reporting effort must be considered. In this study, the degree to which each bird species has been studied both globally and within India was used to account for reporting effort. Reporting effort reassuringly exhibited no association with JEV status among birds, but we nevertheless caution that this finding cannot completely rule out residual bias and will therefore require field-based wildlife surveillance for validation. Second, a number of species were missing data on the life history characteristics, clutch size and egg mass, and especially on population density. Using methods previously validated[25,56] these data were imputed as described in the Methods. Nevertheless, the imputed data do invariably provide a diminished representation of waterbird life history and population density in India, and so future work will benefit from field investigations that can also incorporate life history and population ecology into the monitoring of virus circulation in these birds. Third, the findings are not used to make any specific claims about the true nature and influence of interaction (e.g. competitive exclusion or character displacement) among waterbird communities at various scales because, as previously described, direct observation of interspecific interaction and its effects on community composition was not possible under the framework of the current study. As such, these results will again require validation against field studies of directly observed interaction between bird species at multiple scales. Fourth, despite the exceptionally large number of bird observations obtained for this study, there were insufficient observations for all extant waterbird species in India to model the distribution of each. Therefore, the new estimates presented for species richness and individual species abundances, although the first of their kind for India, are nevertheless based on a sample of bird species (n = 48) and therefore represent a proxy for true species richness and species abundances. Fifth, spatial biases may affect the distribution of species observation data in GBIF due to differential accessibility, as well as the distribution of JEV outbreaks due to differential reporting of cases. Accordingly, background points used in the models were selected proportional to the human footprint and IMR, respectively, to control for these potential biases.

In conclusion, this study has identified optimal preliminary wild waterbird targets for the development of novel One Health JEV surveillance in India. The extent to which the identified species will be validated as important hosts in the infection ecology of JEV, and the specific roles individual species may play (e.g. maintenance vs. amplification hosts), must await the data generated from the implementation of these surveillance mechanisms. Nevertheless, we feel the current results provide the best evidence base to date for actioning new surveillance mechanisms that can effectively couple wildlife, livestock, and human health monitoring at the key sources of interface in landscapes of wetland-rainfed agriculture ecotones.

## Notes

### Competing Interest Statement

The authors have declared no competing interest.

