## Supplemental material for "A preliminary description of the ecological characteristics of wild waterbird Japanese encephalitis virus hosts in high risk landscapes in India"

S1 Table 1. Ardeidae species, Anatidae species, and Rallidae species niche comparisons based on ensemble species distribution models. Each species listed represents that species' modelled suitability with the associated number of observations of the species in the field (and the number of species used for analysis after thinning in parentheses), model fit (deviance), model performance (area under the receiver operating characteristic curve (AUC)), and individual niche overlap with the composite landscape suitability.

| | **Number of field observations** | **Deviance** | **AUC (%)** | **TSS** |
|---|---|---|---|---|
| **Ardeidae species** | | | | |
| *Ardea alba* | 18173 (11794) | 0.83 | 89 | 0.63 |
| *Ardea cinerea* | 17774 (11006) | 0.85 | 89 | 0.62 |
| *Ardea purpurea* | 16406 (9784) | 0.78 | 91 | 0.67 |
| *Ardeola grayii* | 56018 (30755) | 0.87 | 88 | 0.61 |
| *Bubulcus coromandus* | 290 (207) | 0.87 | 88 | 0.64 |
| *Bubulcus ibis* | 53548 (32862) | 0.91 | 87 | 0.58 |
| *Butorides striata* | 3398 (2576) | 0.84 | 89 | 0.64 |
| *Dupetor flavicollis* (*Ixobrychus flavicollis*) | 1602 (1278) | 0.77 | 91 | 0.68 |
| *Egretta garzetta* | 36332 (22326) | 0.87 | 88 | 0.61 |
| *Egretta gularis* | 3081 (2050) | 0.41 | 98 | 0.87 |
| *Egretta intermedia* (*Ardea intermedia*) | 22406 (14473) | 0.82 | 89 | 0.63 |
| *Gorsachius melanolophus* | 112 (107) | 0.54 | 94 | 0.86 |
| *Ixobrychus cinnamomeus* | 2151 (1810) | 0.78 | 91 | 0.67 |
| *Ixobrychus sinensis* | 2300 (1739) | 0.71 | 92 | 0.70 |
| *Nycticorax nycticorax* | 8193 (5473) | 0.86 | 89 | 0.62 |
| **Anatidae species** | | | | |
| *Anas acuta* | 7359 (4975) | 0.86 | 88 | 0.61 |
| *Anas clypeata* | 7832 (5051) | 0.81 | 90 | 0.64 |
| *Anas crecca* | 5760 (3968) | 0.86 | 89 | 0.63 |
| *Anas penelope* | 3566 (2542) | 0.91 | 87 | 0.59 |
| *Anas platyrhynchos* | 1488 (1002) | 0.84 | 89 | 0.62 |
| *Anas querquedula* | 6494 (4430) | 0.83 | 89 | 0.65 |
| *Anas strepera* | 4673 (3133) | 0.85 | 89 | 0.63 |
| *Anser anser* | 2292 (1472) | 0.74 | 92 | 0.68 |
| *Anser indicus* | 4398 (2699) | 0.80 | 90 | 0.66 |
| *Aythya farina* | 3650 (2537) | 0.91 | 87 | 0.60 |
| *Aythya fuligula* | 2241 (1557) | 0.82 | 89 | 0.65 |
| *Aythya nyroca* | 1685 (1184) | 0.71 | 93 | 0.71 |
| *Mergus merganser* | 467 (328) | 0.44 | 97 | 0.86 |
| *Netta rufina* | 2333 (1637) | 0.85 | 89 | 0.62 |
| *Tadorna ferruginea* | 7850 (4986) | 0.86 | 89 | 0.62 |
| *Tadorna tadorna* | 334 (284) | 0.59 | 96 | 0.84 |
| *Anas poecilorhyncha* | 19276 (11654) | 0.82 | 90 | 0.64 |
| *Dendrocygna bicolor* | 854 (581) | 0.45 | 97 | 0.85 |
| *Dendrocygna javanica* | 19493 (11683) | 0.85 | 89 | 0.62 |
| *Nettapus coromandelianus* | 6208 (4220) | 0.82 | 89 | 0.63 |
| *Sarkidiornis melanotos* | 5174 (3401) | 0.80 | 91 | 0.66 |
| **Rallidae species** | | | | |
| *Amaurornis akool* | 879 (755) | 1.04 | 82 | 0.51 |
| *Amaurornis phoenicurus* | 30697 (18558) | 0.87 | 88 | 0.62 |
| *Fulica atra* | 14879 (9367) | 0.87 | 88 | 0.61 |
| *Gallicrex cinerea* | 1539 (1193) | 0.72 | 92 | 0.68 |
| *Gallinula chloropus* | 12284 (7885) | 0.92 | 87 | 0.58 |
| *Gallirallus striatus* | 510 (400) | 0.56 | 96 | 0.81 |
| *Porphyrio porphyrio* | 14860 (8515) | 0.76 | 91 | 0.68 |
| *Porzana bicolor* | 155 (102) | 0.56 | 94 | 0.87 |
| *Porzana fusca* | 1337 (976) | 0.66 | 94 | 0.76 |
| *Porzana pusilla* | 937 (695) | 0.85 | 89 | 0.66 |
| *Rallina eurizonoides* | 145 (123) | 0.69 | 93 | 0.78 |
| *Rallus aquaticus* | 150 (110) | 0.87 | 90 | 0.70 |

S2 Figure 1. Comparison of waterbird species' ecological and life history traits by Japanese encephalitis virus susceptibility status.

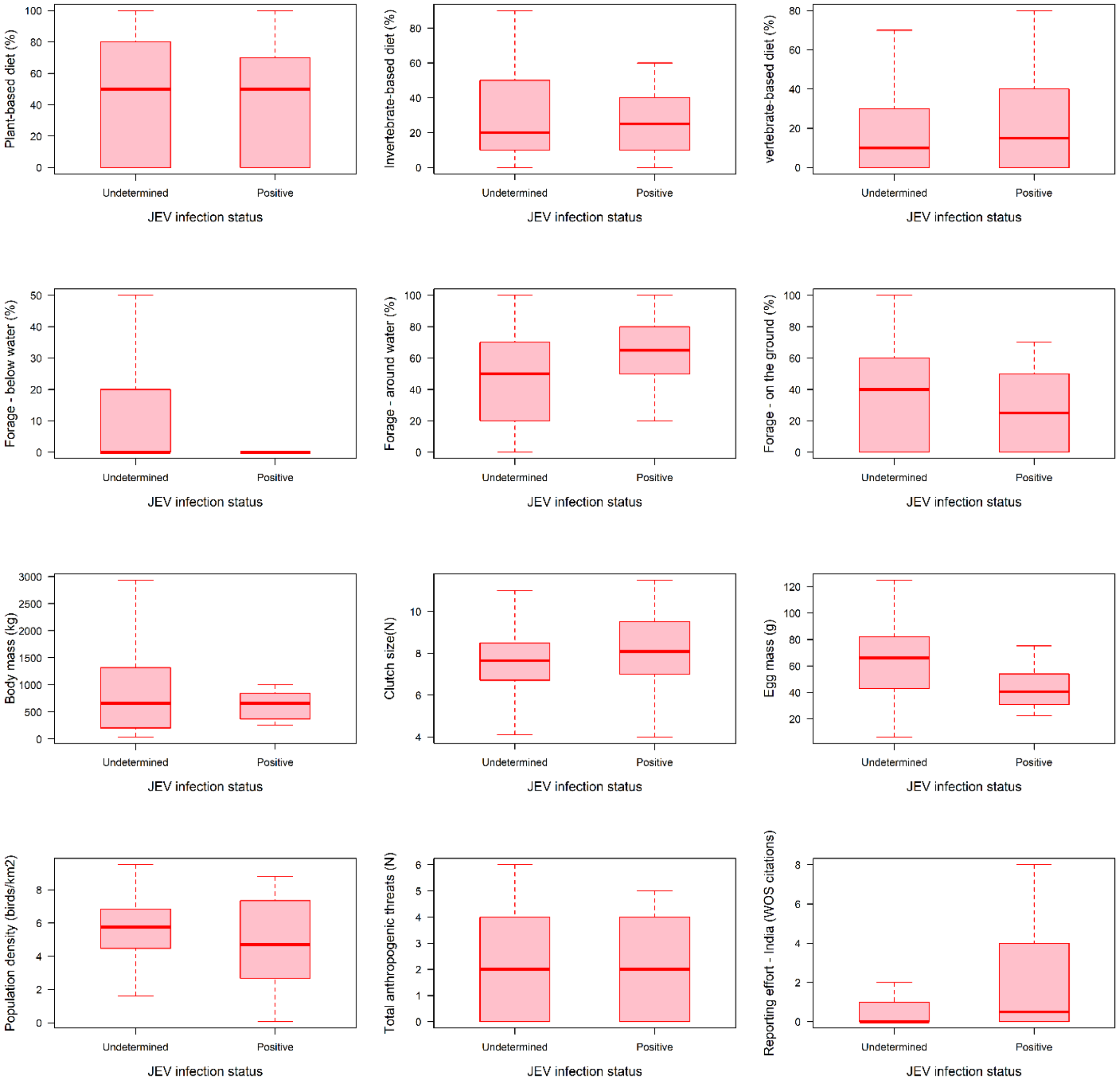

S3 Table 2. Crude bivariate associations between waterbird species traits and Japanese encephalitis virus susceptibility status derived from simple phylogenetic generalised linear models.

| Species trait | Regression coefficient | AIC | p-value |
|---|---|---|---|
| Reporting effort (India) | 0.006 | 97.65 | 0.34 |
| Reporting effort (global) | 0.00002 | 101.52 | 0.80 |
| Body mass (kg) | -0.56 | 97.15 | 0.18 |
| Clutch size (N) | 0.16 | 80.11 | 0.29 |
| Egg mass (g) | 0.003 | 96.89 | 0.39 |
| Increasing population | 1.44 | 79.53 | 0.04 |
| Stable population | -0.31 | 79.53 | 0.71 |
| Threatened due to agriculture | 0.53 | 78.34 | 0.29 |
| Threatened due to urbanisation | 0.71 | 77.98 | 0.23 |
| Threatened due to logging | -1.34 | 79.59 | 0.36 |
| Threatened due to extractive industries | 0.84 | 76.12 | 0.11 |
| Threatened due to road/railway development | 0.18 | 81.68 | 0.83 |
| Threatened due to hunting | -0.0001 | 81.69 | >0.99 |
| Threatened due to invasive species | 0.48 | 79.23 | 0.37 |
| Threatened due to climate change | -0.34 | 80.78 | 0.55 |
| Overall population under threat | -0.001 | 81.77 | >0.99 |
| Diet – plants/seeds | -0.48 | 78.39 | 0.40 |
| Diet – invertebrates | -0.29 | 81.22 | 0.66 |
| Diet – vertebrates | 0.48 | 81.04 | 0.53 |
| Diet – omnivorous | 1.19 | 75.37 | 0.10 |
| IUCN threatened | -1.82 | 76.90 | 0.21 |
| Forage strategy – below water surface | -0.01 | 95.34 | 0.21 |
| Forage strategy – around water surface | 0.03 | 94.31 | 0.02 |
| Forage strategy – on the ground | -0.007 | 96.59 | 0.38 |
| Forage strategy – within the understory | -0.007 | 81.61 | 0.90 |
| Populations density | 0.05 | 77.02 | 0.15 |

S4 Table 3. Regression coefficients and 95% confidence intervals for the associations between Japanese encephalitis virus outbreaks and each species landscape suitability as derived from simple inohomogeneous Poisson point process models. Coefficients represent crude associations from the bivariate models.

| **Bird species landscape suitability** | **Coefficient** | **95% confidence interval** | **AIC** |
|---|---|---|---|
| *Ardea alba* | 0.51 | -0.081 – 1.10 | 573.44 |
| *Ardeola grayii* | 0.42 | -0.170 – 1.00 | 574.35 |
| *Bubulcus coromandus* | **2.56** | **1.867 – 3.254** | **527.08** |
| *Butorides striata* | **2.14** | **1.628 – 2.646** | **512.63** |
| *Egretta garzetta* | 0.21 | -0.384 – 0.798 | 575.78 |
| *Egretta intermedia* | **0.65** | **0.085 – 1.217** | **571.28** |
| *Nycticorax nycticorax* | -0.09 | -0.684 – 0.510 | 576.16 |
| *Anas acuta* | -0.44 | -1.095 – 0.212 | 574.46 |
| *Anas clypeata* | -0.64 | -1.311 – 0.034 | 572.71 |
| *Anas crecca* | **0.72** | **0.047 – 1.398** | **571.90** |
| *Anas Penelope* | **1.12** | **0.498 – 1.745** | **564.05** |
| *Anas platyrhynchos* | **3.91** | **3.359 – 4.457** | **398.29** |
| *Anas strepera* | **1.95** | **1.352 – 2.548** | **535.94** |
| *Aythya fuligula* | **1.72** | **1.166 – 2.282** | **540.59** |
| *Tadorna ferruginea* | **2.56** | **1.906 – 3.221** | **518.35** |
| *Tadorna tadorna* | **2.65** | **2.189 – 3.109** | **460.74** |
| *Anas poecilorhyncha* | **-1.04** | **-1.742 – -0.338** | **567.41** |
| *Dendrocygna bicolor* | **2.73** | **2.316 – 3.146** | **415.32** |
| *Nettapus coromandelianus* | **0.84** | **0.308 – 1.370** | **566.86** |
| *Amaurornis akool* | **2.58** | **1.797 – 3.367** | **533.72** |
| *Fulica atra* | **-1.84** | **-2.570 – -1.099** | **550.58** |
| *Gallinula chloropus* | -0.44 | -1.161 – 0.285 | 574.82 |
| *Rallus aquaticus* | -0.60 | -1.254 – 0.048 | 572.85 |

S5 Table 4. Taluk-level regression coefficients and 95% credible intervals for the associations between Japanese encephalitis virus outbreaks and each species' abundance (per species-present pixel per taluk) as derived from simple integrated nested Laplace approximation models (binomial family). Coefficients represent crude associations from the bivariate models.

| **Bird species abundance/relative abundance** | **Coefficient** | **95% credible interval** | **WAIC** |
|---|---|---|---|
| *Ardea alba* abundance | 0.0007 | -0.0002 – 0.0014 | 569.68 |
| *Ardeola grayii* abundance | 0.0005 | -0.0003 – 0.0012 | 569.95 |
| *Bubulcus ibis coromandus* abundance | **0.0013** | **0.0008 – 0.0019** | **560.04** |
| *Butorides striata* abundance | **0.001** | **0.0007 – 0.0019** | **563.26** |
| *Egretta garzetta* abundance | 0.0006 | -0.0003 – 0.0013 | 569.49 |
| *Egretta intermedia* abundance | **0.0007** | **0.00002 – 0.0014** | **568.01** |
| *Nycticorax nycticorax* abundance | 0.0006 | -0.0005 – 0.002 | 570.03 |
| *Anas acuta* abundance | -0.0014 | -0.004 – 0.0004 | 569.91 |
| *Anas clypeata* abundance | 0.0002 | -0.002 – 0.0014 | 575.78 |
| *Anas crecca* abundance | 0.0004 | -0.0001 – 0.0008 | 570.03 |
| *Anas penelope* abundance | **0.0006** | **0.0001 – 0.0009** | **568.34** |
| *Anas platyrhynchos* abundance | **0.0004** | **0.0002 – 0.0007** | **566.32** |
| *Anas strepera* abundance | **0.0007** | **0.0003 – 0.0010** | **565.53** |
| *Aythya fuligula* abundance | **0.00066** | **0.00026 – 0.0010** | **566.34** |
| *Tadorna ferruginea* abundance | *0.0006* | *0.000 – 0.001* | *618.10* |
| *Tadorna tadorna* abundance | **0.0019** | **0.00095 – 0.0029** | **563.74** |
| *Anas poecilorhyncha* abundance | 0.0006 | -0.0008 – 0.002 | 570.36 |
| *Dendrocygna bicolor* abundance | 0.0027 | -0.0005 – 0.005 | 569.48 |
| *Nettapus coromandelianus* abundance | 0.0006 | -0.00002 – 0.0011 | 570.01 |
| *Amaurornis akool* abundance | **0.0009** | **0.0005 – 0.0013** | **563.21** |
| *Fulica atra* abundance | 0.0000 | -0.0021 – 0.0016 | 572.63 |
| *Gallinula chloropus* abundance | 0.0004 | -0.0004 – 0.001 | 570.73 |
| *Rallus aquaticus* abundance | 0.0005 | -0.0003 – 0.0010 | 570.16 |

S6 Figure 2. Comparison of the distributions of within-family species richness, species abundances, relative species abundances, at the community level (taluk) after adjusting for the biotic constraints of sympatry. Maps are used only for the purposes of species richness and do not reflect the authors' assertion of territory or borders of any sovereign country including India.

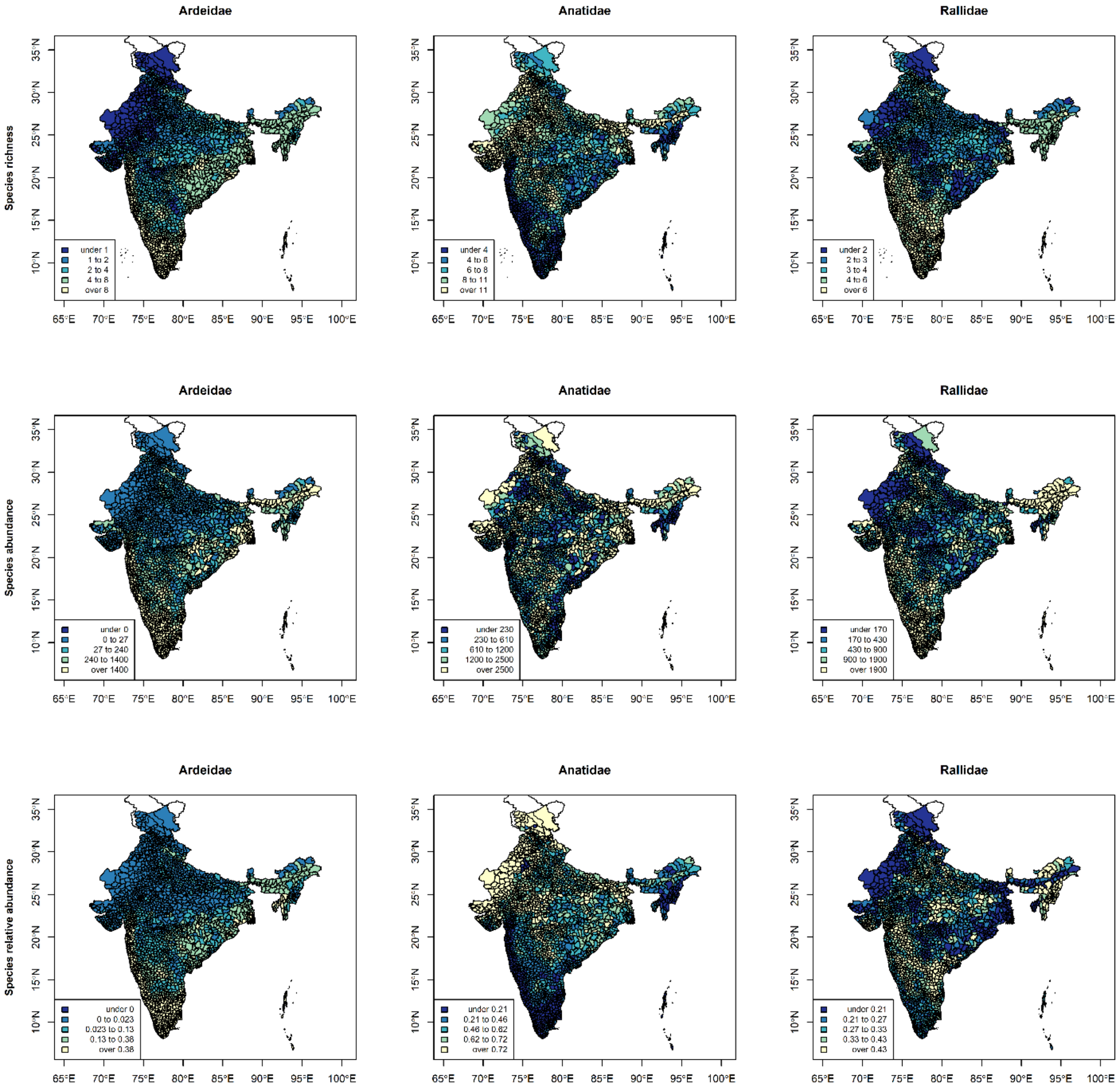
